# Agriculturally developed areas reduce genetic connectivity for a keystone neotropical ungulate

**DOI:** 10.1101/2024.01.22.576460

**Authors:** Mozart Sávio Pires Baptista, Alexine Keuroghlian, Leandro Reverberi Tambosi, Marina Corrêa Côrtes, Fernanda de Góes Maciel, Douglas William Cirino, Gabriela Shmaedecke, Cibele Biondo

## Abstract

Modified landscapes can restrict the movement of organisms, leading to isolation and reduced population viability, particularly for species with extensive home ranges and long-distance travel, such as white-lipped peccaries (WLPs, *Tayassu pecari*). Previous studies have indicated that forested areas favor WLP herd movements, but the impact of the non-forested areas on their genetic connectivity is unknown. In this study, we used land cover, the Brazilian Roads Map, and population genetic data to investigate the impact of non-forested matrices on WLP’s genetic connectivity in the Pantanal floodplain and surrounding Cerrado plateau of central-west Brazil. We compared isolation-by-distance (IBD), isolation-by-barrier, and isolation-by-resistance models and tested 39 hypotheses within a modeling framework. Finally, we identified the optimal areas for ecological corridors based on the most effective landscape model. Barrier and landscape resistance were more strongly correlated with genetic relatedness than the IBD model. The model that received the most robust support considered only forest as habitat. All other classes formed a matrix that impeded gene flow, including agriculture, grassland, savannah, and paved and unpaved roads. WLP herds living in landscapes with reduced forest cover are more vulnerable to the effects of genetic isolation. To maintain gene flow, it is essential to establish connections between habitats throughout the landscape. Conservation programs should prioritize strategies that strengthen connections between habitats, including facilitating wildlife road-crossing structures and creating/restoring ecological corridors to link isolated habitat fragments.

## 1. INTRODUCTION

Land use and land cover (LULC) transformations drive global environmental change, significantly contributing to the ongoing loss of biodiversity (Foley et al., 2005). These changes primarily result in habitat loss and fragmentation, reducing landscape connectivity (Edelsparre et al., 2018). This leads to population fragmentation, threatening species’ survival chances (Fahrig, 2003). This threat is further amplified by the increasing genetic differentiation between isolated populations and the decrease in genetic variability within populations (Fauvergue et al., 2012; Willi et al., 2013). Eventually, the population fitness will decline, and the risk of local extinction becomes greater (Hoffmann et al., 2017).

The field of landscape genetics examines how LULC’s changes impact microevolutionary processes (Manel et al., 2003). One approach employs isolation models, such as Isolation by Resistance (IBR) and Isolation by Barrier (IBB), to infer connectivity patterns. IBR considers the permeability of the matrix (McRae, 2006), whereas IBB considers abrupt disruptions caused by barriers such as roads or rivers (Cushman et al., 2006). The variation in estimates of pairwise genetic distances explained by IBR or IBB is compared to the variation explained by geographic distances alone, referred to as Isolation by Distance models (IBD; Wright, 1943). IBD is considered the simplest landscape genetics pattern and is expected to occur without landscape effects (Balkenhol et al., 2009; Jenkins et al., 2010). However, the contribution of IBR and IBB to highly mobile social species is not yet well understood, particularly in the Neotropics (Monteiro et al., 2019). Considering the significant human-mediated landscape changes in the Neotropics (Rull, 2020), it is crucial to identify landscape features that reduce gene flow between populations to improve landscape management and conservation programs.

White-lipped peccaries (WLPs, *Tayassu pecari*) are the only Neotropical ungulates that form large herds, sometimes with up to 300 individuals, which periodically split into smaller sub-herds based on resource availability (Keuroghlian et al., 2004; Keuroghlian et al., 2022). These social animals inhabit vast areas of native forests, ranging from 2000 to 20000 ha, and they can move long distances, up to 10 km/day (Keuroghlian et al. 2004; Keuroghlian & Eaton, 2008; Keuroghlian et al., 2009; Beck et al., 2017). Although relatedness is associated with their social behavior (sociogenetic structure - Rufo, 2012; Nóbrega, 2018), there is high gene flow between herds, resulting in weak spatial genetic structure, even in relatively well-conserved large areas, where we expect some genetic structure by distance (with approximately 30% of individuals in herds being migrants - Biondo et al., 2011; Baptista et al., in prep.).

An analysis of WLP populations across Brazil, including Cerrado, Atlantic Forest, and Pantanal areas, reveals that isolation by distance occurs at distances greater than 180 km (Maciel et al., 2019). It is known that the presence of an agricultural matrix can impede the movement of herds at a landscape scale, requiring well-connected habitats for viable populations to persist (Jorge et al., 2020). A study in the Atlantic Forest revealed significant genetic differentiation between WLP populations living in relatively close forest fragments (10 and 36 km apart). This differentiation was attributed to the hindrance of herd movement by the agricultural matrix (Martin, 2018). The impact of LULC changes on gene flow between historically connected populations remains unclear. Therefore, it is crucial to investigate the impact of agricultural areas on the dispersal of WLPs to ensure the conservation of their populations and the protection of their habitats. To achieve this goal, it is essential to develop strategies that promote the preservation and restoration of well-connected landscapes.

Road construction significantly impacts gene flow as it modifies the surrounding habitat, creating less favorable conditions for wildlife. These alterations result in habitat degradation, reduced forest cover, and limited resource availability (Jaeger et al., 2005). Consequently, specialized animals often display road avoidance behavior, which can result in long-term genetic isolation (Ceia-Hasse et al., 2018). For instance, WLPs are closely linked to forest habitats and predominantly rely on a frugivorous diet. Approximately 80% and 60% of their diet consisted of native fruits in the Atlantic Forest and Pantanal, respectively (Desbiez et al. 2009, Keuroghlian et al. 2009, Keuroghlian & Eaton 2008, Keuroghlian, Eaton, and Desbiez, 2009). Therefore, expecting the population to survive in areas lacking high-quality resources and availability, such as those near roads, is unrealistic.

Several studies support the hypothesis that WLPs tend to avoid roads. Jorge et al. (2013) observed a lower probability of encountering WLP herds near roads. Moreover, documented cases of WLP roadkill were infrequent (Abra et al., 2020). However, when roadkill incidents occur, they can have catastrophic consequences, resulting in the simultaneous death of multiple animals within the herd. A study considering the species’ entire distribution area estimated that over 4,000 individuals are killed by road accidents annually in Brazil, without accounting for avoidance behavior, population density, or local extinctions (Pinto et al., 2022). Moreover, WLP has been observed to avoid conventional wildlife crossing tunnels beneath roadways (Abra et al., 2020). GPS tracking data further revealed that while WLP may cross unpaved or dirt roads (depending on the traffic), they strongly avoid crossing paved roads (Oshima, 2019). Based on these studies, paved roads are expected to act as significant barriers to gene flow among WLP herds, while unpaved or dirt roads may be less restrictive. Nonetheless, further research is necessary to comprehensively assess the impact of roads as barriers to gene flow among WLP populations.

WLPs have experienced population declines and local extinctions throughout their distribution range (Altrichter et al., 2012; Keuroghlian et al., 2012), mainly due to habitat loss and growing human activity (Reyna-Hurtado et al., 2010; 2016). As a result, the species is currently classified globally as vulnerable (IUCN, 2020). Understanding how land-use changes impact genetic connectivity in WLP populations is crucial for conservation. In this study, we used land-use maps and genetic data to assess the impact of landscape features on WLP genetic relatedness in the Brazilian Pantanal basin and surrounding plateau. We aimed to 1) examine how different land-use types affect WLP gene flow, 2) assess the barrier effect of paved and unpaved roads, and 3) identify optimal ecological corridors. We hypothesized that agricultural lands cause isolation by resistance (IBR), while paved roads lead to isolation by barrier (IBB). We expected a barrier effect from paved roads but not unpaved roads, given the higher human influence near paved roads (Lupinetti-Cunha et al., 2022; Tisler et al., 2022). Finally, we considered isolation by distance (IBD) as a plausible model only if neither of these landscape elements had a substantial effect.

## 2. MATERIAL AND METHODS

### 2.1. STUDY AREA AND SAMPLING

Our study was conducted in the Upper Paraguay Basin (UPB) in Mato Grosso do Sul, Brazil, encompassing both the Pantanal floodplain and highlands (plateau). The Pantanal, the world’s largest contiguous wetland, features grasslands, savannas, and forests shaped by annual flood cycles (Pott et al., 2011; Junk et al., 2014). The highlands consist mainly of Cerrado savanna, recognized as a biodiversity hotspot due to its rich species diversity and species at risk (Myers et al., 2000). This region experiences a distinct seasonal tropical savanna climate characterized by wet summers and dry winters (Köppen-Geiger, Aw). The Pantanal floodplain in our areas comprises 28% forests and 72% savannas, aquatic grasslands, and pasture/monoculture areas (Eaton et al., 2017). The highlands alternate between savanna fragments and seasonal forests, surrounded by pastures and monocultures (Jorge et al., 2019).

We collected samples from 179 adult individuals at 51 sampling points across nine localities (Figure 1). We followed the capture and blood sampling methods described by Maciel et al. (2019). To avoid spatial autocorrelation, we grouped sampling points within a 12 km radius, exceeding the genetic neighborhood size of approximately 6.5 km (Baptista et al., in prep.). Thus, we analyzed nine localities representing sampling units spaced from 12.6 to 138.1 km apart (Appendix S1).

**Figure 1.**
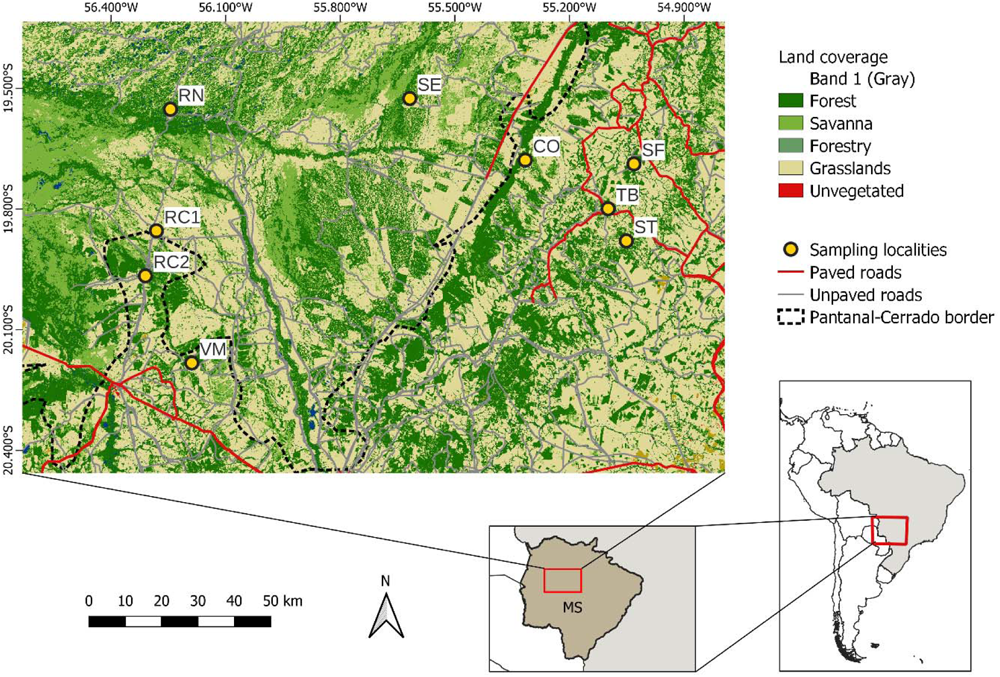
Study area with the distribution of the nine sampling localities in the Pantanal plain and the adjacent plateau in Mato Grosso do Sul, Brazil. RN = Rio Negro Farm, SE = Santa Emília Farm, RC1 = Refúgio Ecológico Caiman (Pantanal), RC2 = Refúgio Ecológico Caiman (Plateau), VM = Vinte e Três de Março Farm, CO = Colorado Farm, ST = Santa Tereza Farm, SF = Safira Farm, and TB = Taboco. Source of land use and land cover mapping: MapBiomas 4.0 (2018).

### 2.2. DATASET

We genotyped samples using 13 microsatellite loci, as outlined by Maciel et al. (2019). Their analysis confirmed that the herds under study constituted a single genetic population, with a low FST value of 0.03 ± 0.01. Microsatellite markers were chosen for their effectiveness in identifying subtle genetic structures on a fine spatial scale, even in recently altered landscapes (Anderson et al., 2010). All analyzed loci were polymorphic, with 2 to 13 alleles (mean: 5.92 ± 3.88), and observed and expected heterozygosities averaged 0.52 ± 0.23 and 0.58 ± 0.20, respectively. Four loci deviated from Hardy-Weinberg equilibrium (p < 0.02; IGF1, SW857, Tpec5, and Tpec10), with evidence of null alleles in two (IGF and Tpec5).

To explore genetic isolation models, we used an individual-based approach, calculating the coefficient of relatedness as described by Wagner et al. (2006). This coefficient quantifies genetic similarity based on the probability of sharing identical alleles by common descent, informing genetic relationships, population structure, gene flow, and consanguinity. We computed mean relatedness between individuals within localities to obtain a single measure for each sampling point. The coefficient of relatedness accommodated null alleles and was computed using the ML-RELATE program (Kalinowski et al., 2006).

For landscape variables, we utilized Land Cover and Use Maps from the MapBiomas 4.0 historical series (mapbiomas.org), which provides data on Brazil’s land cover and use. The series was generated through automated classification using satellite imagery and machine learning techniques. We incorporated the Brazilian Roads Map, including paved and unpaved roads data (Lupinetti-Cunha et al., 2022).

### 2.3. LANDSCAPE DATA ANALYSIS

We computed landscape distances at a 30 m spatial resolution. Given the stability of land use and land cover (LULC) classes over the past three decades (Appendix S2), we utilized data from 2006, coinciding with the primary sample collection period. Our analysis encompassed a total area of 150,000 km², including a 20,000 km² buffer around sampling points. LULC classes covering less than 1% were omitted. We amalgamated native and exotic grasslands into a single “grasslands” class due to their structural similarity. In the case of forests, we consolidated rivers and all native forests into a single category, recognizing riparian forests as crucial movement areas for the species (Keuroghlian & Eaton, 2008) with minimal resistance. The “agriculture” class was based on the cultivation of rice, coffee, wheat, corn, beans, cassava, cotton, peanuts, sugarcane, and predominantly soybeans. Consequently, for the genetic-landscape correlation, we considered six classes: 1) forests, 2) savannas, 3) grasslands, 4) agriculture, 5) unpaved roads, and 6) paved roads.

### 2.4. PARAMETERIZATION OF THE LANDSCAPE RESISTANCE

Landscape resistance analysis involves creating a hypothetical resistance surface using a land use and land cover (LULC) map, assigning cost values to each class to represent organism movement. Various methods exist for determining these cost values (Zeller, McGarigal, & Whiteley, 2012).

To evaluate LULC-imposed resistance, we examined 12 alternative landscape scenarios (Figure 2) by assessing the impact of different combinations of LULC classes and roads. These scenarios fell into three groups: The first, a null model, followed the isolation by distance (IBD) model (Figure 2A), where all LULC classes were equally permeable to animal movement (Wright, 1943). The second group represented the Isolation by Resistance (IBR) model (Figure 2 D-L), which considered both favorable (habitat) and resistant (matrix) areas for dispersion (McRae, 2006). The third group comprised the Isolation by Barrier (IBB) model, which extended the IBD (Figure 2 B-C) and IBR models (Figure E, F, H, I, K, L) to account for roads as potential barriers to movement (Cushman et al., 2006). The IBB models were split into two categories: IBBp (considering only paved roads) and IBBup (considering all roads, paved or unpaved). In testing the hypotheses of IBBp and IBBup, we assigned a fixed road resistance value of 200, exceeding the maximum resistance (100) (Figure 2, Appendix S3).

**Figure 2.**
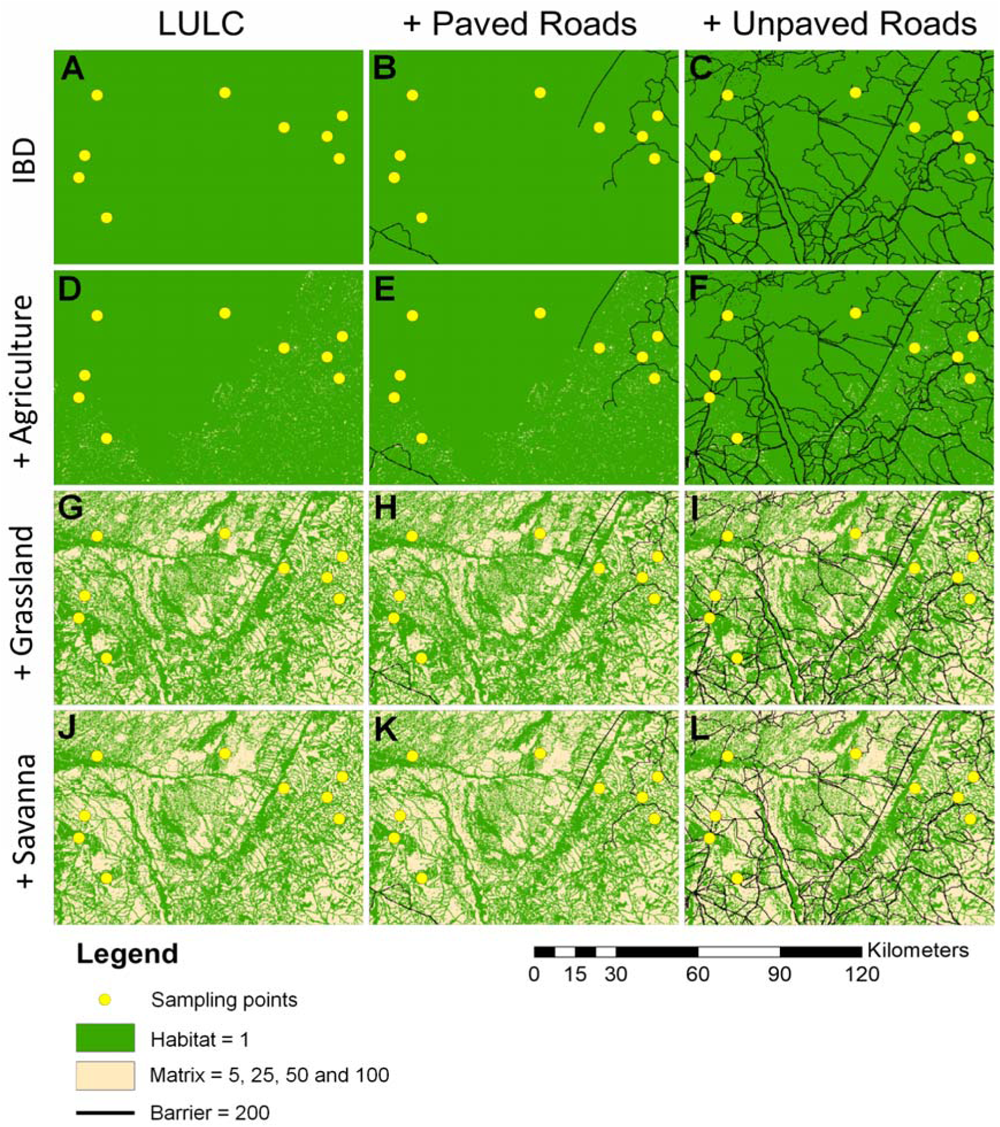
Landscape surfaces used to evaluate Isolation by Distance (A), Isolation by Resistance (D-L), and Isolation by Barrier hypothesis (B, C, E, F, H, I, K, L). Land use and land cover (LULC) classes were divided into two permeability classes (habitat and matrix), with the lowest permeability class including as matrices: agriculture only (D, E, F), agriculture plus grassland (G, H, I), and agriculture plus grasslands plus savannas (J, K, L), in these last models just forest is considered habitat. Isolation by Barrier models considered paved roads (B, E, H, K) or all roads (C, F, I, L).

To identify the best-fit scenario explaining WLP genetic connectivity, we allocated distinct cost values to LULC classes, adopting a parametrization approach: setting habitat areas to 1, matrix areas to x-values (5, 25, 50, or 100), and roads to 200. We selected these predetermined values based on prior research recommendations (Shirk et al., 2010; Cushman et al., 2013). Lower x-values denoted lower movement costs, while higher x-values indicated higher costs (Ruiz-Gonzales et al., 2014). Consequently, the tested surfaces could have up to three cost values: all LULC equally permeable (habitat, IBD); LULC categorized as habitat or matrix (two cost values); LULC considered as habitat, matrix, or barrier (three cost values). Under these parametrizations, we evaluated 39 landscape permeability hypotheses (Appendix S4) by calculating effective resistance distances between WLP individuals for each hypothetical surface. We accounted for multiple paths over multiple generations using Electrical Circuit Theory (McRae, 2006; McRae & Beier, 2007; McRae et al., 2008) implemented in Circuitscape 4.0 (McRae et al., 2013). Circuitscape identifies the most permeable paths, considering various possible routes, yielding an optimized surface with paired resistance distance values.

### 2.5. LANDSCAPE GENETICS ANALYSIS

To identify the best genetic isolation model given the alternative resistance surfaces, we correlated the relatedness coefficient with the geographic distance (IBD) – the null hypothesis – or resistance distances from each model (IBR or IBB) based on Mantel tests (Mantel, 1967) and partial Mantel tests (Smouse et al., 1986) using a three-step approach. All tests were implemented in the package “Ecodist” (Goslee & Urban, 2007) in R version 4.0.2 (R Development Core Team, 2020) with 10,000 permutations. In the first step, we ranked the best-supported candidate model using the maximum simple Mantel correlation between pairwise relatedness and landscape resistance distances, i.e., the higher the correlation, the more supported the model (Cushman & Languth, 2010). We then use an original (second step) and reciprocal (third step) causal modeling approach to reduce errors associated with incorrect correlations and due to the impossibility of applying a model selection criterion (Cushman et al., 2006; 2013).

### 2.6. ORIGINAL CAUSAL MODELING

The original causal modeling framework determined the best genetic isolation models (Cushman et al., 2006). In this step, the alternative models (IBR and IBB) were tested against the null model (IBD), as described in Cushman et al. (2006). We performed two sets of partial Mantel tests: (1) partial Mantel tests between relatedness and landscape resistance distances, partializing the effects of geographical distance; and (2) partial Mantel tests between relatedness and geographical distance, partializing the effects of landscape resistance distances. Because distance matrices from the landscape are being contrasted with a matrix of genetic similarity (relatedness coefficients), we expected that if there is an effect of the landscape on genetic isolation, the values of simple Mantel tests and partial Mantel tests should be (1) positive and significant, and the values of (2) should be negative or not significant (Cushman et al., 2006).

## 3. RESULTS

The Simple Mantel tests found a positive correlation between genetic relatedness and geographic distance, the IBD model (r = 0.87, p = 0.001). However, all other simple Mantel tests showed a higher Pearson correlation coefficient than the IBD model, indicating a stronger association between genetic relatedness and the tested landscape models (Figure 3). As the complexity of the landscape models increased, so did the correlation between landscape resistance distances and genetic relatedness. Among the tested models, the L100 model exhibited the highest correlation (r = 0.999; p = 0.001; Figure 3, Appendix S4). The L100 model considers the forest class as habitat (low-cost value) and the land classes agriculture, grassland, and savannah as highly resistant (with a cost value of 100). Also, paved and unpaved roads imposed a barrier effect, with a resistance value of 200.

**Figure 3.**
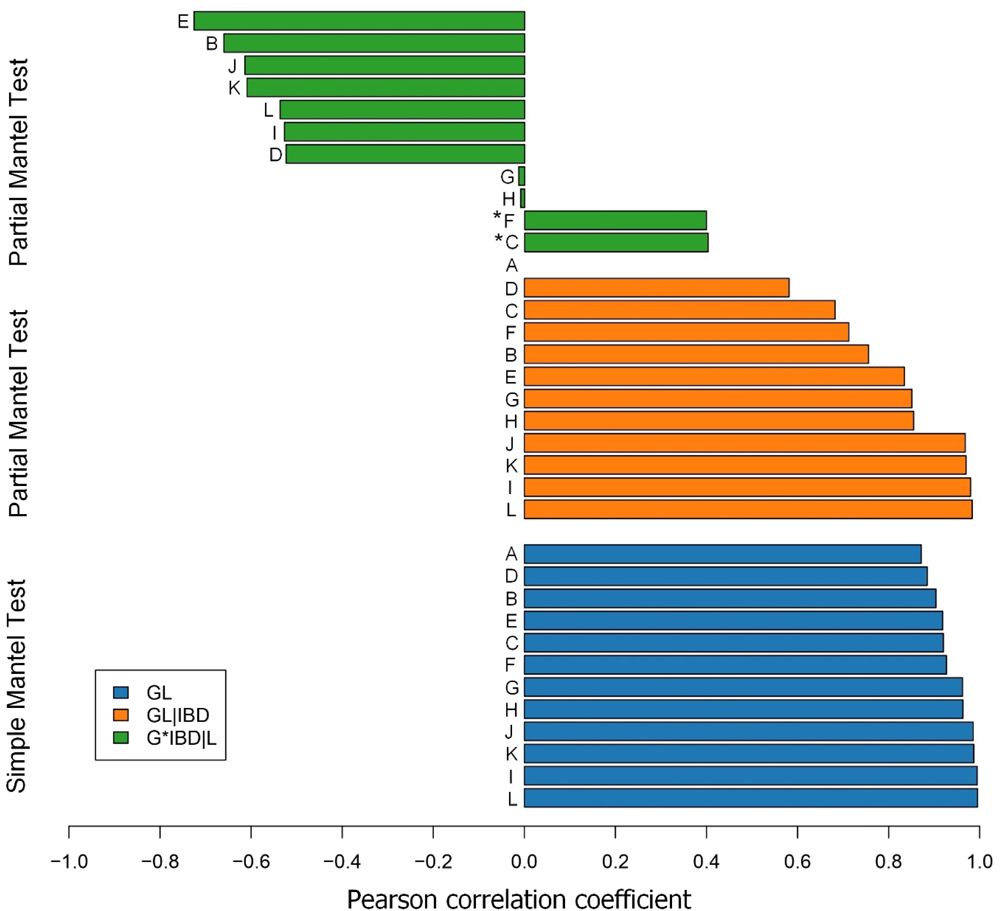
Ranking of best correlation models between genetic (G) and landscape (L) data, resulting from original causal modeling, with three types of Mantel tests: blue bars represent simple Mantel tests (G*L), orange bars represent partial Mantel tests that partialize the effect of isolation by distance (G*L|IBD), and green bars represent partial Mantel tests that partialize the effect of the tested model (G*IBD|L). The resistance models are denoted as follows: A = Isolation by distance (IBD), B = A + paved roads, C = B + unpaved roads, D = A + agriculture, E = D + paved roads, F = E + unpaved roads, G = D + grassland, H = G + paved roads, I = H + unpaved roads, J = G + savannah, K = J + paved roads, L = K + unpaved roads. The values represent the mean of resistance values assigned to LULC classes in each model (5, 25, 50, and 100). Non-significant models (p > 0,05) are denoted with *.

In the original causal modeling, the L100 model remained superior when the IBD effect was removed in the partial Mantel tests (R = 0.997, p = 0.001; Figure 3). Furthermore, when the effect of L100 was removed from the IBD model, the IBD model was not significant (R = - 0.559, p = 1, Appendix S4).

Reciprocal causal modeling demonstrated that only L100 (Model 39) maintained a positive correlation among all models when the effects of all other models were removed. Also, it was the only model that caused a negative correlation in all other models when its effect was removed (Appendix S5). Therefore, our analyses suggest that the L100 model is the most plausible explanation for the effect of landscape on WLP’s genetic relatedness.

Finally, we used the L100 model to identify landscape areas with potential for ecological corridors (Figure 4) based on the highest resistance values (red) and permeability values (blue). We observed that there are low-cost pathways connecting most areas. However, VM and SE areas appear to be more isolated. We also noticed that the two main existing corridors of native vegetation are in areas with lower electrical currents, and there are pixels of low cost that coincide with the presence of roads.

**Figure 4.**
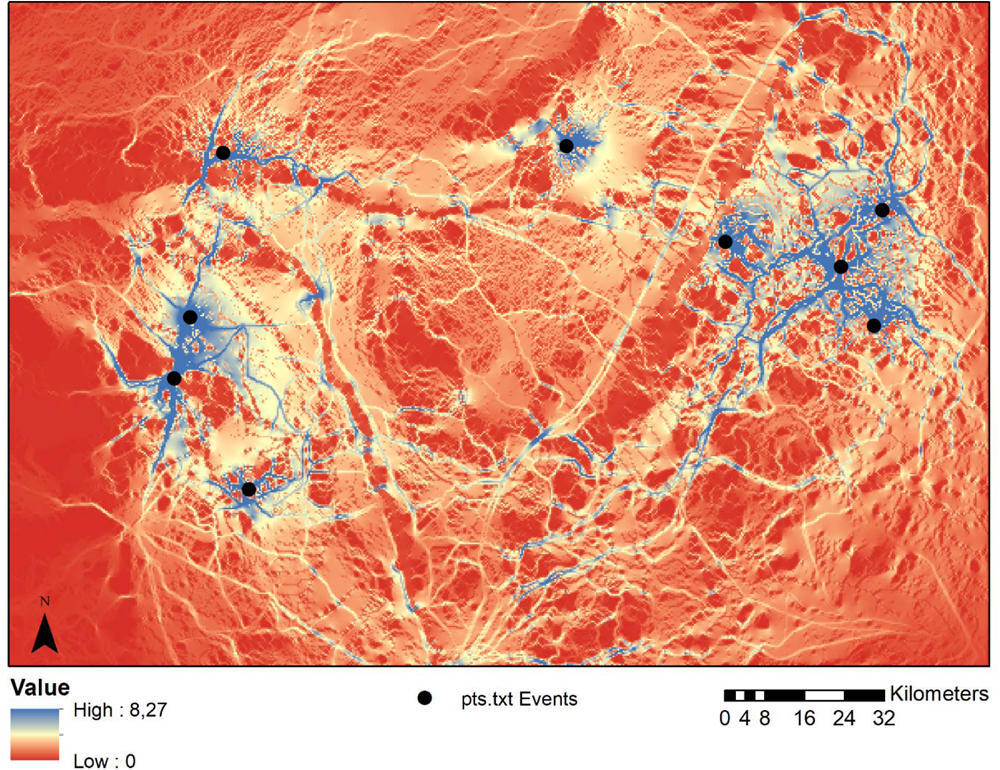
Map of the landscape parameters that affect the movements of white-lipped peccaries (WLP). The L100 model, consisting of forested habitat cover and non-forested land cover matrices (including savannas, grasslands, and agriculture) and linear barriers (paved and unpaved roads), was the best supported. Black circles represent the sampling localities. The red cells on the map indicate areas with the highest cost to WLP movement, while the blue cells indicate areas with the lowest costs.

## 4. DISCUSSION

In our study, we delved into landscape features influencing gene flow in highly mobile animals with extensive home ranges, particularly focusing on White-lipped Peccaries (WLPs) inhabiting Brazilian wetlands and savannas. We aimed to determine how non-forested areas affect gene flow among WLP herds, potentially leading to reduced genetic relatedness between individuals in regions with low forest connectivity.

Initially, we employed the IBD, IBB, and IBR models to analyze genetic relatedness patterns in the landscape. While IBD showed a significant effect, we enhanced our model’s predictive accuracy by incorporating landscape variables into the resistance surface. The most effective model (L100) designated forested areas as suitable habitats, whereas other land cover types like agriculture, grassland, and savanna were deemed less favorable, posing higher costs to WLP movement. Our analysis pinpointed that both paved and unpaved roads acted as substantial barriers to WLP gene flow within our study area. These findings underscore the urgency of conserving well-connected landscapes to mitigate human activities’ detrimental effects on WLP populations. Moreover, our study illuminated that even native vegetation areas, such as savannas, can exhibit resistance to gene flow, with secondary unpaved roads posing similar challenges to paved ones, emphasizing the significance of roadless areas for WLP conservation.

Previous studies have demonstrated that land use and cover types, such as agriculture, pasture, and urban areas, can reduce gene flow and genetic diversity, with roads being particularly detrimental (Bowman et al., 2010; Rytwinski & Fahrig, 2012; Miles et al., 2019). Species like the WLP that actively avoid roads may be reducing the risk of roadkill, but in doing so, they also increase the risk of long-term genetic isolation (Holderegger & Di Giulio, 2010; Teixeira et al., 2020). The presence of roads has been widely reported to reduce genetic diversity in other ungulate populations (Ito et al., 2013; D’Amico et al., 2016). Our findings align with this, indicating that both paved and unpaved roads constitute significant genetic barriers for WLPs. In our study area, unpaved roads may even pose a greater threat due to their higher abundance, frequent heavy vehicle use, and potential for habitat alteration and runoff (Trombulak & Frissell, 2000). Such consequences are significant for WLP populations and their long-term viability.

While our study revealed cost-effective pathways indicating substantial connectivity, some regions, notably VM and SE, exhibited lower connectivity, posing challenges for WLP movement. Native vegetation corridors in high-cost regions may not be as effective as anticipated. Low-cost pixels coinciding with roads are concerning, as roads contribute to habitat loss and mortality. Conservation strategies should prioritize promoting connectivity and gene flow between isolated habitats. Establishing ecological corridors, stepping stones, and permeable matrices can facilitate habitat connection and organismal movement (Rudnick et al., 2012; de la Fuente et al., 2018; Huang et al., 2018). It is essential to identify the reasons behind corridor failures and prioritize alternative land use practices like sustainable agriculture and biodiversity-focused land management.

The resistance map generated in our study serves as a foundation for future research, informing conservation and management strategies to enhance ecological connectivity and resilience in fragmented landscapes. This map enables the evaluation of strategies aimed at promoting gene flow and improving the long-term viability of wildlife populations, crucial for maintaining ecological balance in fragmented landscapes.

The Biodiversity Impact Reduction Plan for Land Roads (PRIM-IVT) in Brazil assessed road impacts differently in the Pantanal and surrounding plateau (Falcon et al., 2018). Although plateau populations face high highway-related effects, Pantanal WLPs experience lower to moderate impacts, with unpaved roads offsetting the perception that paved roads are more restrictive. Our research found no significant difference between these road types. Roads significantly harm WLP populations, but they are not a priority in PRIM-IVT, despite their “Vulnerable” status according to IUCN and ICMBio. We recommend designating WLPs as a priority in PRIM-IVT and implementing tailored strategies, including corridors, to mitigate road impacts.

To mitigate road-related impacts on wildlife populations, effective solutions are essential to enhance habitat connectivity. Strategies include establishing ecological corridors and species-specific faunal passages, especially in the Pantanal, where well-designed faunal passages within alluvial forests can cater to natural wildlife movement. The effectiveness of such structures should be thoroughly evaluated, possibly incorporating passage sections at the base of bridges or enhancing existing structures. These measures can significantly improve connectivity and the long-term viability of wildlife populations affected by road fragmentation, fostering a harmonious coexistence between road infrastructure and biodiversity.

While our cost values were based on existing literature, it’s important to acknowledge that they may only partially reflect reality. Future studies should explore alternative parameterization methods, such as ResistanceGA (Peterman, 2018), or incorporate biological movement data from GPS tracking or radio necklace technology, to develop more accurate resistance models reflecting landscape fragmentation costs (Elliot et al., 2014; Unnithan Kumar et al., 2022). These efforts are crucial for mitigating the negative impacts of landscape fragmentation on wildlife populations.

In conclusion, our study underscores the pivotal role of landscape resistance in shaping WLP genetic connectivity in fragmented landscapes. Our findings enhance our understanding of WLP movement behavior and identify priority paths for ecological corridor planning. However, we also highlight significant challenges in conserving the species, such as genetic structure and habitat connectivity issues. Comprehensive conservation strategies prioritizing habitat connectivity and ecological corridor planning are essential. Our easily replicable approach can provide valuable insights to decision-makers involved in conservation planning for WLPs and other threatened species in fragmented landscapes.

## Acknowledgment

We thank Tatiana P. Freitas, Ezidio Arruda, Celso Vicente, Maria do Carmo A. Santos, Paulino A. O. Santos, Renata R. Rocha, Julia E. de F. Oshima, and Maria L. da S. P. Jorge for their assistance with field work. This work was funded by the Fundação de Amparo à Pesquisa do Estado de São Paulo - FAPESP (grant number 2015/20133-0), Conselho Nacional de Desenvolvimento Científico e Tecnológico - CNPq (grant number 479760/2012-8), Fundação Manoel Barros, Earthwatch Institute volunteers, CI-Brasil, Global Ecotours & Expeditions volunteers, Silicon Valley Community Foundation, The Overbrook Foundation, and WCS Brasil. We also thank the staff and landowners from Caiman Ecological Refuge, Vinte e Três de Março Farm, Rio Negro Farm, Santa Emilia and Campo Lourdes Farm; and landowners in the Municipality of Corguinho. This work was carried out with the support of the Coordenação de Aperfeiçoamento de Pessoal de Nível Superior - Brazil (CAPES) - Financing Code 001.

